# Spectral graph theory of brain oscillations – revisited and improved

**DOI:** 10.1101/2021.09.28.462078

**Authors:** Parul Verma, Srikantan Nagarajan, Ashish Raj

## Abstract

Mathematical modeling of the relationship between the functional activity and the structural wiring of the brain has largely been undertaken using non-linear and biophysically detailed mathematical models with regionally varying parameters. While this approach provides us a rich repertoire of multistable dynamics that can be displayed by the brain, it is computationally demanding. Moreover, although neuronal dynamics at the microscopic level are nonlinear and chaotic, it is unclear if such detailed nonlinear models are required to capture the emergent meso- (regional population ensemble) and macroscale (whole brain) behavior, which is largely deterministic and reproducible across individuals. Indeed, recent modeling effort based on spectral graph theory has shown that an analytical model without regionally varying parameters can capture the empirical magnetoencephalography frequency spectra and the spatial patterns of the alpha and beta frequency bands accurately.

In this work, we demonstrate an improved hierarchical, linearized, and analytic spectral graph theory-based model that can capture the frequency spectra obtained from magnetoencephalography recordings of resting healthy subjects. We reformulated the spectral graph theory model in line with classical neural mass models, therefore providing more biologically interpretable parameters, especially at the local scale. We demonstrated that this model performs better than the original model when comparing the spectral correlation of modeled frequency spectra and that obtained from the magnetoencephalography recordings. This model also performs equally well in predicting the spatial patterns of the empirical alpha and beta frequency bands.

**Highlights:**

- We show an improved hierarchical, linearized, and analytic spectral graph theory-based model that can capture the frequency spectra obtained from magnetoencephalography recordings
- This model also accurately captures the spatial patterns of the empirical alpha and beta frequency bands, requiring only 5-10 graph eigenmodes to do so

## 1 Introduction

How the brain exhibits dynamic functional activity with a static anatomical structural wiring, and how the structural and functional alterations are associated with neurological diseases [1] has been a long-standing open question in the field of neuroscience. Functional activity in the gray matter regions can be measured non-invasively using functional magnetic resonance imaging (fMRI), electroencephalography (EEG), and magnetoencephalography (MEG). The connections between the gray matter regions, or the structural wiring, is estimated using diffusion tensor imaging (DTI) from MRI. One way of investigating the relationship between the functional activity and the structural wiring is by exploring a macroscopic graph with the gray matter regions as the nodes and the weights of the edges determined by the extent of connectivity of the white matter fibers between the nodes. Subsequently, various graph theoretic and modeling approaches have been undertaken to determine the functional-structural relationships and how they are altered in various neurological diseases.

Graph theoretic approaches have been widely used to evaluate statistical measures of relationships between functional and structural connectivity [2–12]. However, they do not take into account any details of neural physiology [13] and therefore are limited in their capability to develop a mechanistic understanding.

On the other hand, detailed computational modeling approaches have been undertaken to incorporate neural activity, such as neural mass [14, 15] and mean field modeling [16–18], where neural assemblies and connections among them are modeled. Such computational models are non-linear and require several time consuming simulations to obtain the structure-function relationships. Although neuronal dynamics at the microscopic level (single neurons) are nonlinear and chaotic, it is unclear if such detailed nonlinear models are required to capture the emergent meso- (regional population ensemble) and macro-scale (whole brain) behavior, which is largely deterministic and reproducible across individuals [19, 20]. Indeed, it has been suggested that brain-wide neural activity can be independent of microscopic local activity of individual neurons [21, 22, 17, 6, 23, 13], and instead may be regulated by long-range structural connectivity [24–27]. Based on this hypothesis, Raj and colleagues had earlier developed a hierarchical, linear, analytic spectral graph theory model (SGM) which could accurately capture empirical MEG spectra and spatial distribution of alpha and beta frequency bands [28].

SGM provides a closed-form solution of brain oscillations in the form of steady state frequency response obtained from the eigen-decomposition of a graph Laplacian, based on spectral graph theory [29–32]. This model incorporates the structural connectome and long-range axonal conduction delays, and provides a set of frequency-rich spectra which can be directly compared to the empirical MEG wideband spectra obtained after source reconstruction of MEG signals. Moreover, since the analytical solution to this SGM model can be obtained in a closed-form, no time-consuming simulations are required. The model is parameterized by *a small set of global parameters:* the neural gains, time constants, conduction velocity, and macroscopic coupling. These model parameters admit physical interpretations and can be potentially controlled by neuromodulation [33]. Most importantly for downstream applications, SGM model inference is highly feasible in comparison to other methods, and it is the only model whose inference can be realized from empirical regional power spectra rather than from functional correlation structures like functional connectivity (FC). Finally, the spectral graph model is conceptually attractive because it places the *graph* explicitly at the center of the model, a counter-point to coupled Neural Mass Models (NMMs), where the effect of the graph comes about only indirectly as coupling coefficients between local neural masses.

The previously published SGM had local model elements derived from a control theory viewpoint, due to which the local parameters and gain terms lacked classical interpretability in terms of extant NMMs. Here, we reformulate the SGM local model from bottom-up, in line with classical neural masses. We thoroughly explore the inference of this modified model on real MEG data, and show that the modified model is essentially equivalent to the original SGM, and displays a small but significantly more favorable fit to real MEG regional power spectra. Thus, the new model keeps the key strengths of the original, while providing more biologically interpretable parameters, especially at the local scale. This modified SGM (M-SGM) has an excellent ability to capture the spatial patterns of empirical alpha and beta frequency bands, requiring only 5-10 graph eigenmodes to do so.

## 2 Methods

### 2.1 M-SGM

We hierarchically model the local cortical mesoscopic and long-range macroscopic signals for each brain region, where the regions are obtained using the Desikan-Killiany atlas [34]. We then solve the model equations to obtain a closed-form solution in the Fourier frequency domain. This provides us frequency rich spectra which is an estimate of the source-reconstructed MEG spectra.

**Notation** All the vectors and matrices are written in boldface and the scalars are written in normal font. The frequency *f* of a signal is specified in Hertz and the corresponding angular frequency *ω* = 2*πf* is used to obtain the Fourier transforms. The connectivity matrix is defined as ***C*** = *c_jk_*, where *c_jk_* is the connectivity strength between regions *j* and *k*, normalized by the row degree.

#### 2.1.1 Mesoscopic model

For every region *k* out of the total *N* regions, we model the local excitatory signal *E_k_*, local inhibitory signal *I_k_* as well as the long range excitatory signal *x_k_* where the global connections are incorporated. The local signals are modeled using an analytical and linearized form of neural mass equations. We write a set of differential equations for evolution of *E_k_* and *I_k_* due to decay of individual signals with a fixed neural gain, incoming signals from populations that alternate between excitatory and inhibitory signals, and input white Gaussian noise. Letting *f_e_*(*t*) and *f_i_*(*t*) denote the ensemble average neural impulse response functions, the equations for *E_k_* and *I_k_* are:

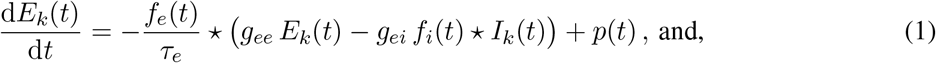

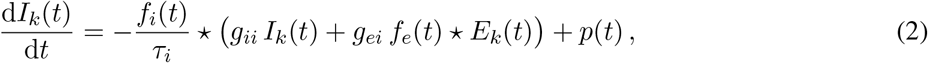

where, * stands for convolution, *p*(*t*) is input noise, parameters *g_ee_, g_ii_, g_ei_* are neural gain terms, and parameters *τ_e_, τ_i_* are characteristic time constants. These are global parameters and are the same for every region *k*.

Here, the ensemble average neural impulse response functions *f_e_*(*t*) and *f_i_*(*t*) are assumed to be Gamma-shaped and written as the following:

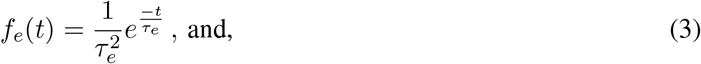

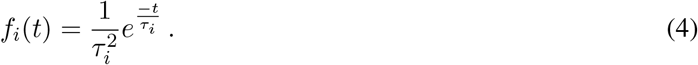

#### 2.1.2 Macroscopic model

A similar equation is written for the macroscopic signal *x_k_*, for every *k^th^* region, accounting for long-range excitatory corticocortical connections for the pyramidal cells. The evolution of *x_k_* is assumed as a sum of decay due to individual signals with a fixed excitatory neural gain, incoming signals from all other connected regions determined by the white matter connections, and the input signal *E_k_* (*t*) + *I_k_* (*t*) determined from Eq. (1), (2). The equation for *x_k_* is the following:

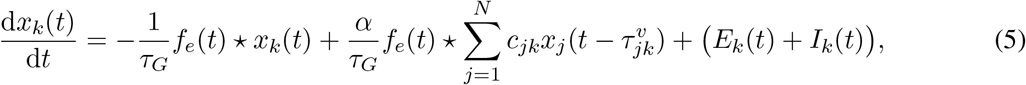

where, *τ_G_* is the graph characteristic time constant, *α* is the global coupling constant, *c_jk_* are elements of the connectivity matrix determined from DTI followed by tractography, 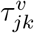 is the delay in signals reaching from the *j^th^* to the *k^th^* region, *υ* is the cortico-cortical fiber conduction speed with which the signals are transmitted. The delay 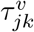 is calculated as *d_jk_/υ*, where *d_jk_* is the distance between regions *j* and *k*.

These set of equations are parameterized by 8 global parameters: excitatory time constant *τ_e_*, inhibitory time constant *τ_i_*, macroscopic graph time constant *τ_G_*, excitatory neural gain *g_ee_*, inhibitory neural gain *g_ii_*, alternating population neural gain *g_ei_*, global couping constant *α*, and conduction speed *υ*. The neural gain *g_ee_* is kept as 1 to ensure parameter identifiability. We estimate the 7 global parameters using an optimization procedure described in the next section.

#### 2.1.3 Model solution in the Fourier domain

Since the above equations are linear, we can obtain a closed-form solution in the Fourier domain as demonstrated below. The Fourier transform 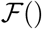 is taken at angular frequency *ω* which is equal to 2*πf*, where *f* is the frequency in Hertz. The Fourier transform of the equations (1), (2) will be the following, where 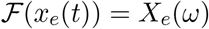 and 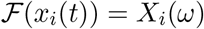:

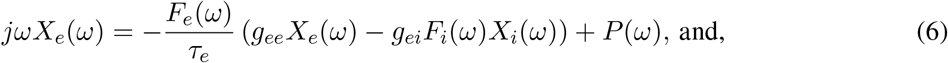

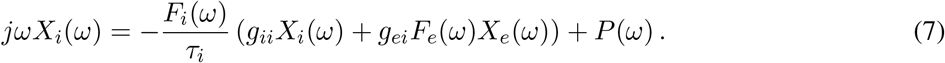

Here, *P*(*ω*) is the Fourier transform of the input Gaussian noise *p*(*t*) which we assume to be identically distributed for both the excitatory and inhibitory local populations for each region, and the Fourier transforms of the ensemble average neural response functions are the following:

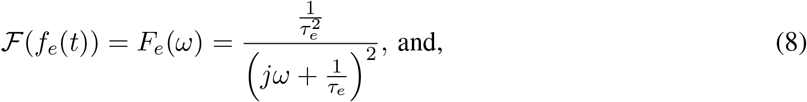

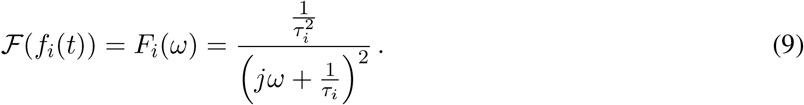

On solving the above equations (6), (7), we get the following expressions for *X_e_*(*ω*) and *X_i_*(*ω*):

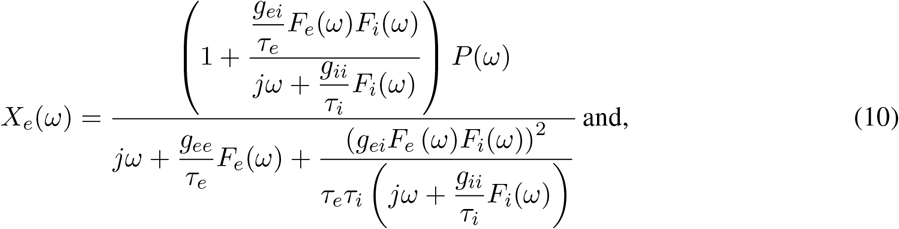

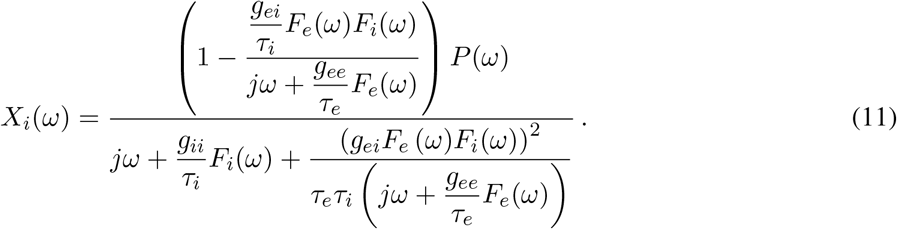

Then, the transfer functions *H_e_*(*ω*) and *H_i_*(*ω*) can be separated out and we can write the following:

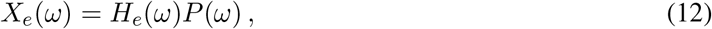

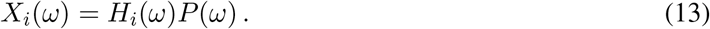

The total neural population can be written as the following:

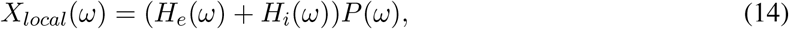

thus, *H_local_*(*ω*) = *H_e_*(*ω*) + *H_i_*(*ω*).

In order to obtain a Fourier response of the macroscopic signal, we first re-write Eq. (5) in the vector form as the following:

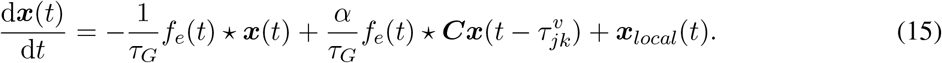

We similarly take a Fourier response of the macroscopic signal and obtain the following as the Fourier transform of Eq. (15), where 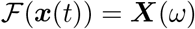:

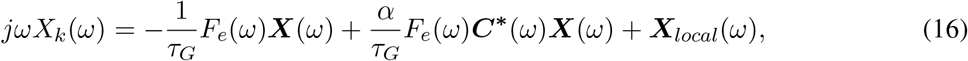

where, 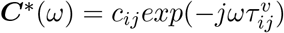. Note that each element in the matrix *C* is normalized already by the row degree. The above equation can be re-arranged as the following:

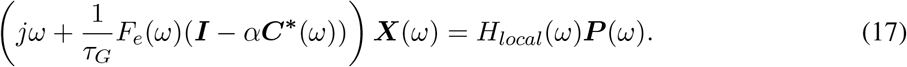

Here, we define the complex Laplacian matrix:

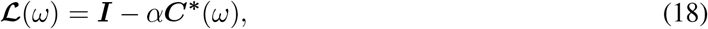

where, I is the identity matrix of size *NXN*. The eigen-decomposition of this complex Laplacian matrix can be written as:

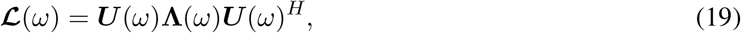

where, ***U***(*ω*) are the eigenvectors and **Λ**(*ω*) = diag([λ_1_(*ω*),…, λ*_N_*(*ω*)]) consist of the eigenvalues λ_1_(*ω*),…, λ*_N_*(*ω*), at angular frequency ω. The macroscopic response Eq. (17) can be written as:

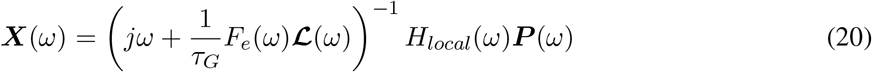

By using the eigen-decomposition of the Laplacian matrix, the above equation can be re-written as the following:

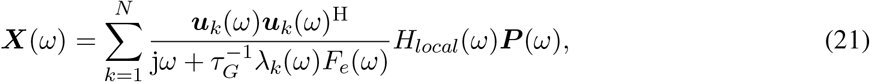

where, ***u**_k_*(*ω*) are the eigenvectors from ***U***(*ω*) and λ*_k_*(*ω*) are the eigenvalues from **Λ**(*ω*) obtained by the eigen-decomposition of the complex Laplacian matrix 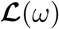 obtained in Eq. 19. Equation (21) is the closed-form steady state solution of the macroscopic signals at a specific angular frequency *ω*. We use this modeled spectra to compare against empirical MEG spectra and subsequently estimate model parameters.

### 2.2 Model parameter estimation

The dataset used for this work is the same as the one used for the previous SGM work [35]. For this dataset, MEG, anatomical MRI, and diffusion MRI was collected for 36 healthy adult subjects (23 males, 13 females; 26 left-handed, 10 right-handed; mean age 21.75 years, age range 7-51 years). Data collection procedure has already been described previously [28]. All study procedures were approved by the institutional review board at the University of California at San Francisco and are in accordance with the ethics standards of the Helsinki Declaration of 1975 as revised in 2008. MEG recordings were collected while the subjects were resting and had eyes closed. MRI followed by tractography was used to generate the connectivity and distance matrices. MEG recordings were downsampled to 600 Hz, followed by a bandpass filtering of the signals between 2 to 45 Hz using `firls` in MATLAB [36] and generation of the frequency spectra for every region of interest using the pmtm algorithm in MATLAB [36].

Modeled spectra was converted into power spectral density (PSD) by calculating the absolute value of the frequency response and both modeled and empirical MEG spectra were converted to to dB scale by taking 20log_10_() of the PSD. Pearson’s r between modeled PSD and the empirical PSD was used a goodness of fit metric for estimating model parameters. Pearson’s r was calculated for comparing spectra for each of the regions, and then they were averaged over all the 68 cortical regions. This average Pearson’s r was the objective function for optimization and used for estimating the model parameters. We used a dual annealing optimization procedure in Python and performed parameter optimization both for the M-SGM and the original SGM [37]. Parameter initial guesses and bounds are specified in Table 1. The bounds were the same as that in the original study. The dual annealing optimization was performed for three different initial guesses, and the parameter set leading to maximum Pearson’s correlation coefficient was chosen for each subject. The dual annealing settings were: `maxiter = 500`. All the other settings were the same as default.

**Table 1:** M-SGM parameter values, initial guesses, and bounds for parameter estimation

| Name | Symbol | Initial value 1 | Initial value 2 | Initial value 3 | Lower/upper bound for optimization |
| --- | --- | --- | --- | --- | --- |
| Excitatory time constant | $\tau_e$ | 0.012 s | 0.018 s | 0.006 s | [0.005 s, 0.02 s] |
| Inhibitory time constant | $\tau_i$ | 0.005 s | 0.01 s | 0.018 s | [0.005 s, 0.02 s] |
| Long-range connectivity coupling constant | $\alpha$ | 1 | 0.5 | 0.1 | [0.1, 1] |
| Transmission velocity | $v$ | 5 m/s | 10 m/s | 18 m/s | [5 m/s, 20 m/s] |
| Alternating population gain | $g_{ei}$ | 4 | 2 | 1 | [0.5, 5] |
| Inhibitory gain | $g_{ii}$ | 1 | 2 | 4 | [0.5, 5] |
| Graph time constant | $\tau_G$ | 0.006 s | 0.01 s | 0.018 s | [0.005 s, 0.02 s] |
| Excitatory gain | $g_{ee}$ | n/a | n/a | n/a | n/a |

## 3 Results

### 3.1 M-SGM fits MEG spectra better

The local mesoscopic model specified in equations (1), (2) is a modification of the original SGM. Here, we have introduced asymmetry in the two equations by including the term for alternating populations. In the original SGM, the equations for excitatory and inhibitory signals were identical and the alternating population’s transfer function was modeled separately. Therefore, we compare the performance of the two models. We estimated parameters for both the models and then compared the modeled spectra. In particular, we compared the Pearson’s r between the empirical MEG spectra and the modeled spectra in both the cases for the same set of subjects, as shown in Fig. 1A. M-SGM has a statistically significantly higher Pearson’s r than SGM, based on a paired t-test of the Fisher’s z-transformed Pearson’s r. Moreover, Pearson’s r was higher when M-SGM was used instead of SGM for almost every subject, as shown by the blue lines connecting subject-wise points corresponding to the two models.

**Figure 1:**
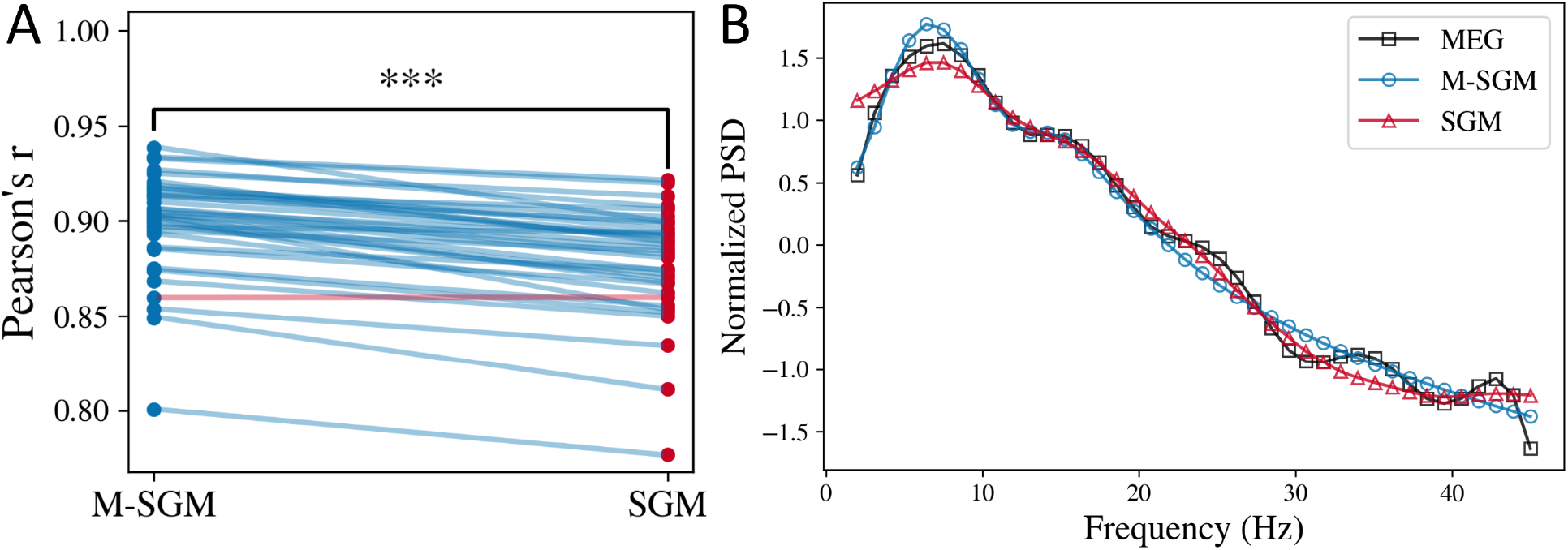
Modified SGM fits MEG spectra better. **A:** Comparison of average Pearson’s r for the modified versus the original SGM. Each line is joining the Pearson’s r corresponding to a specific subject. The line is in blue color if M-SGM’s Pearson’s r is higher than SGM’s Pearson’s r for that subject, and it is in red for vice versa. In this plot, each line other than one is in blue, implying for most subjects, M-SGM’s Pearson’s r was higher than SGM’s Pearson’s r. **B:** Average normalized power spectral density (PSD) obtained from MEG, M-SGM, and original SGM.

As seen in Fig. 1B, the average normalized power spectral density predicted by both the original and the modified SGM fit well with the empirical MEG spectra recorded. Both the models can exhibit a primary alpha peak and a secondary beta peak, however, they cannot generate the smaller peaks observed in the MEG spectra at higher frequencies. It is also to be noted that we only estimate the shape of the spectra, not the scale. Therefore, we used Pearson’s r as a metric for the goodness of fit. The scale can be adjusted by scaling the noise term *P*(*ω*) appropriately, which will be a part of the future work.

### 3.2 Both M-SGM and SGM predict alpha and beta spatial distributions

We also wanted to investigate if the eigenmodes 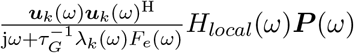 (for every *k*) can capture the spatial distribution of the empirical alpha and beta frequency bands, as was shown for the original SGM. To this end, we sorted the modeled spectral eigenmodes according to their spatial correlation with the empirical alpha and beta bands spatial distribution, respectively. Here, we generated the spatial distribution of alpha and beta bands by summing the MEG spectra from frequencies 8-12 Hz and 13-25 Hz, respectively. The spatial correlation was defined as the Pearson’s r between the regional distribution of the summed alpha and beta spatial distribution and the model predicted eigenmodes which were also summed within the respective frequency bands. After sorting the eigenmodes, we summed the first few eigenmodes that lead to a peak in the spatial correlation. We used these sorted and summed eigenmodes as predictors of the alpha and the beta band activity.

Figure 2 and 3 demonstrates the spatial correlation of the eigenmodes. As seen in Figs. 2B and 3B, the spatial correlation peaks with a subset of eigenmodes and then decreases again, and is similar for both the M-SGM and the SGM. The black curve in both the cases represent the average of the spatial correlations. Fig. 2A demonstrates the regional distribution of the empirical MEG spectra in the cortical region and that predicted by M-SGM, specifically its sum of top eigenmodes in the alpha band, for two different subjects whose spatial correlations are shaded in blue and brown in Fig. 2B, respectively. The Pearson’s r for Fig. 2A are 0.66 and 0.62 for the left and right columns, respectively. The modeled alpha power spatial distribution is prominent in the posterior regions, which strongly accords with the MEG literature, where alpha peak is well-known to be strongly posterior during rest and eyes closed situations. Based on a one-sided paired t-test between the maximum Pearson’s r obtained from M-SGM and those obtained from SGM, there was no significant difference, with a p-value of 0.799. Both M-SGM and SGM performed similarly in predicting the alpha frequency band’s spatial distribution.

**Figure 2:**
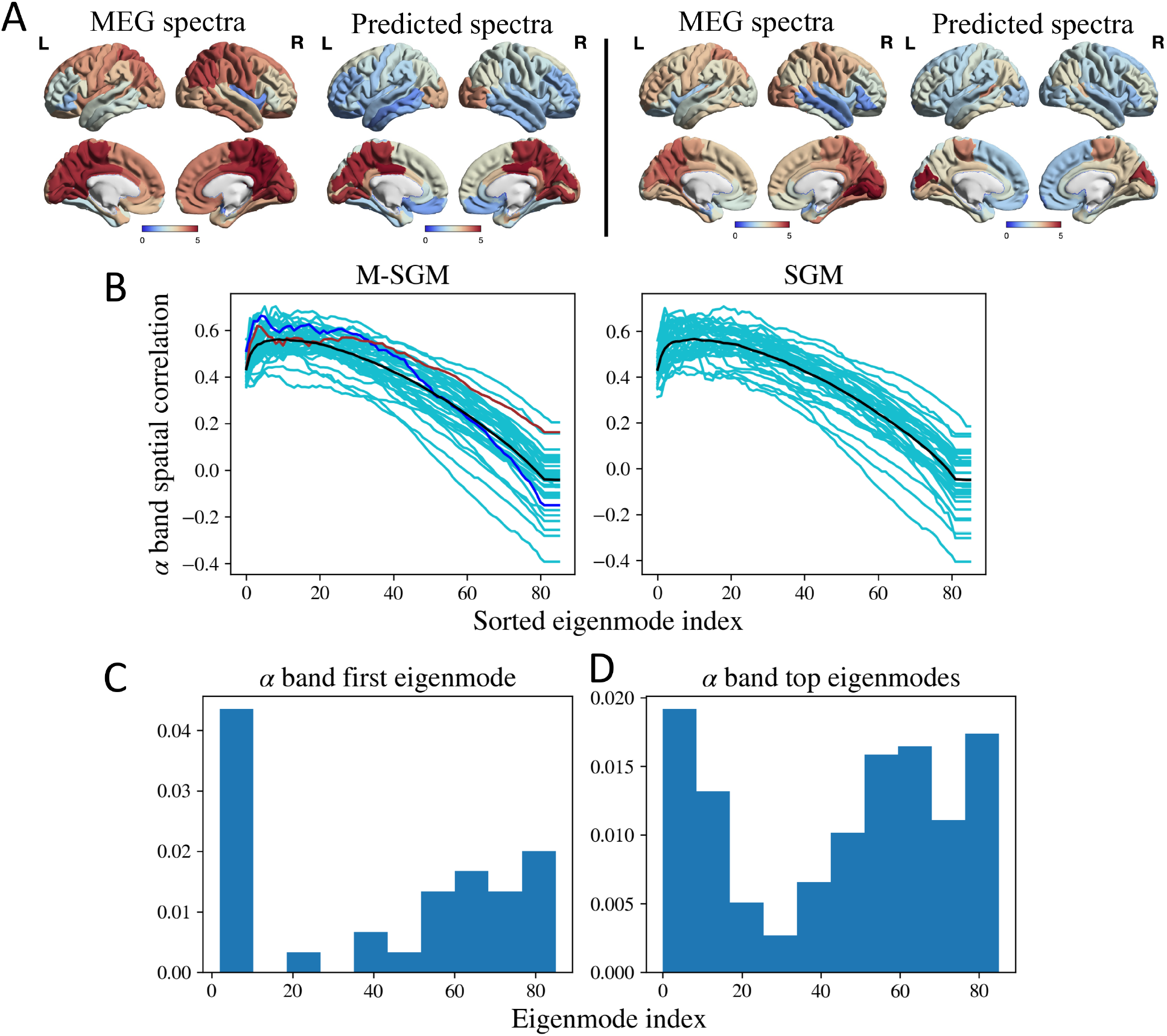
**A:** Left: MEG spectra and sum of top eigenmodes in the alpha band for a specific subject, shaded in blue in B. The Pearson’s r was 0.66. Right: MEG spectra and sum of top eigenmodes for a different subject, shaded in brown in B. The Pearson’s r was 0.62. **B:** Spatial correlations for alpha frequency band. The cyan lines correspond to subject-specific spatial correlations obtained as more eigenmodes are added. The black line corresponds to the average of the spatial correlations. The blue and brown lines correspond to two subjects whose spatial distribution has been demonstrated in A. P-value based on a one-sided paired t-test of the maximum Pearson’s r between M-SGM and SGM gave *p* = 0.799. No statistically significant difference was found between the two. **C:** Histogram of first eigenmode with maximum spatial correlation with the alpha frequency band. **D:** Histogram of the top eigenmodes whose summation leads to maximum spatial correlation with the alpha frequency band. Brain surface renderings were generated using BrainNet Viewer [38] in MATLAB [36].

**Figure 3:**
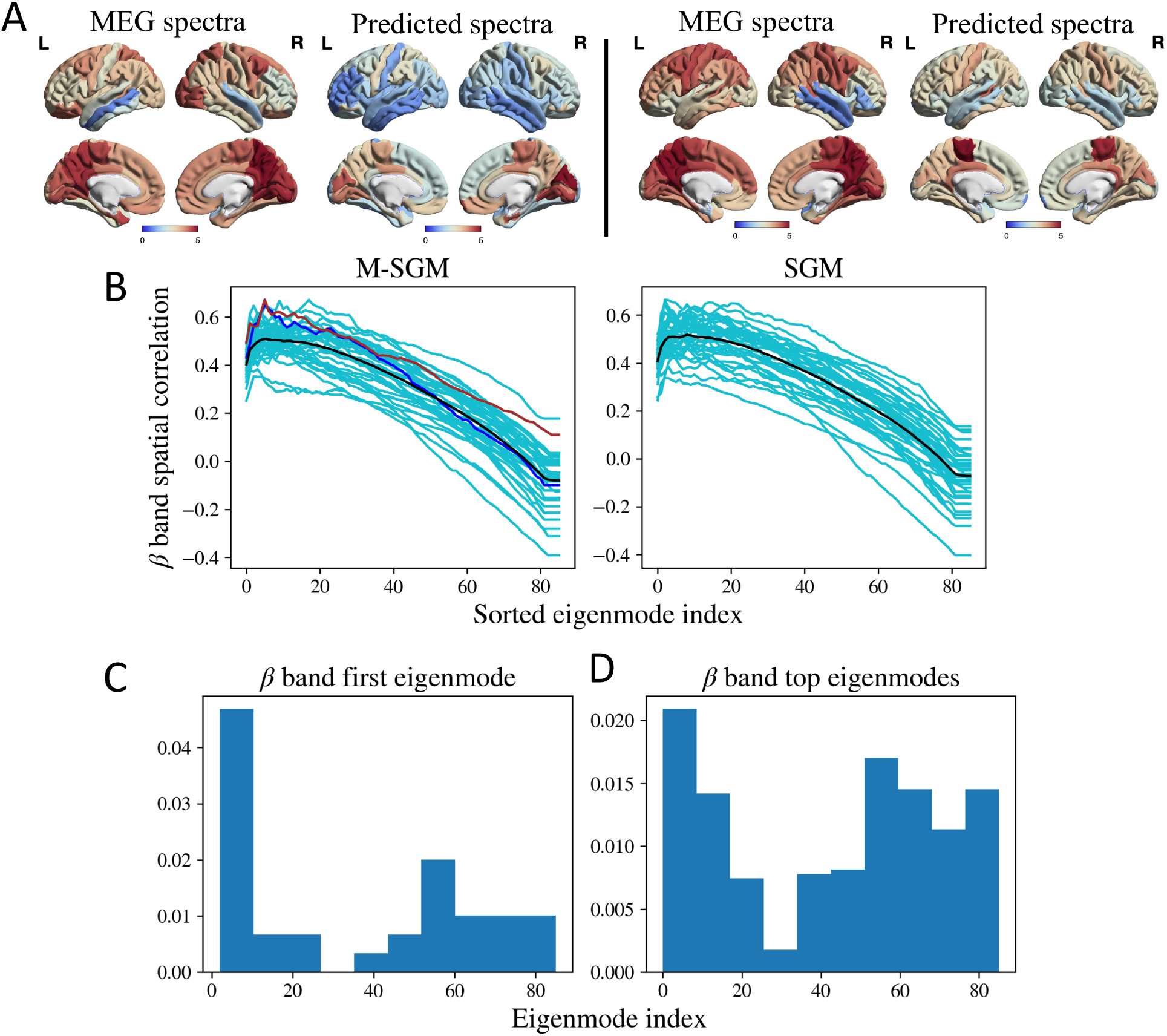
**A:** Left: MEG spectra and sum of top eigenmodes in the beta band for a specific subject, shaded in blue in B. The Pearson’s r was 0.65. Right: MEG spectra and sum of top eigenmodes for a different subject, shaded in brown in B. The Pearson’s r was 0.67. **B:** Spatial correlations for beta frequency band. The cyan lines correspond to subject-specific spatial correlations obtained as more eigenmodes are added. The black line corresponds to the average of the spatial correlations. The blue and brown lines correspond to two subjects whose spatial distribution has been demonstrated in A. P-value based on a one-sided paired t-test of the maximum Pearson’s r between M-SGM and SGM gave *p* = 0.873. The maximum Pearons’s r of M-SGM is statistically not different from that of SGM for the beta band. **C:** Histogram of first eigenmode with maximum spatial correlation with the beta frequency band. **D:** Histogram of the top eigenmodes whose summation leads to maximum spatial correlation with the beta frequency band. Brain surface renderings were generated using BrainNet Viewer [38] in MATLAB [36].

Fig. 3A demonstrates the regional distribution of the empirical MEG spectra and sum of top eigenmodes in the beta band, for two different subjects whose spatial correlations are shaded in blue and brown in Fig. 3B, respectively. The Pearson’s r for Fig. 3A are 0.65 and 0.67 for the left and right columns, respectively. Based on a one-sided paired t-test between the maximum Pearson’s r obtained from M-SGM versus those obtained from SGM, no statistically significant difference was found between the two, with a *p*-value of 0.873. Both M-SGM and SGM performed similarly in predicting the beta frequency band’s spatial distribution.

In order to explore which graph eigenmodes most contribute to the summed model in above figures, we plot in Fig. 2C and D and Fig. 3C and D histograms of the eigenmode index of each participating eigenmode. We show both the single most-maximally contributing eigenmode, as well as all eigenmodes that appear up to the peak observed in Figs. 2B and 3B. We also observed that neither the first eigenmode with maximum spatial correlation, nor the top eigenmodes whose sum lead to the maximum spatial correlation, are necessarily those that appear as the first few eigenmodes as ordered by their eigenvalue.

## 4 Discussions

In this work, we demonstrated that a biophysical linearized spectral graph model can generate frequency-rich spectra that accurately match the empirical MEG spectra, with an overall accuracy higher than that of the original SGM. SGM contained local model elements derived from a control theory viewpoint, due to which the local parameters and gain terms lacked classical interpretability in terms of extant neural mass models. The current reformulation brings the M-SGM in line with classical neural masses. In M-SGM, we have introduced an asymmetry in the excitatory and inhibitory population equations due to the alternating populations term. This asymmetry is the key distinguishing feature between M-SGM and SGM, and is the reason why the M-SGM is biophysically more realistic.

This modified model retains all the attractive features of the original SGM: it is hierarchical, analytic, graph-based, and is parameterized by a parsimonious set of biophysically interpretable global parameters: the neural gains, time constants, conduction velocity, and macroscopic coupling. These model parameters admit physical interpretations and can be potentially controlled by neuromodulation [33]. Due to its closed-form analytical solution in terms of frequency responses, the M-SGM can be fitted directly to regional MEG power spectra, without requiring any time-consuming simulations or numerical integration.

Due to their parsimony a key feature of this and the original SGM is that parameter inference is far more tractable than various non-linear modeling approaches such as The Virtual Brain [39], where multiple refinement steps are required for inference of a larger set of free parameters. While SGM and M-SGM cannot yield a repertoire of dynamical solutions that the detailed non-linear modeling counterparts can, it is unclear if incorporating such non-linearities are required at the macroscopic scale [20]. In comparison to methods based on coupled NMMs, the inference of SGM and M-SGM model parameters inference can be realized directly from empirical wideband regional power spectra rather than from functional correlation structures like FC. Indeed, cutting edge tools like The Virtual Brain do not attempt to fit to empirical wideband power spectra, relying instead on capturing the second-order correlation structures of brain activity after filtering out high-frequency signals or removing the carrier frequency and retaining only the (slowly-varying) Hilbert amplitude envelope. The latter aspect was analyzed elegantly using a linearized coupled NMM by Tewarie et al. [40]; however their model is only accessible via numerical integration and time series simulations, unlike ours.

M-SGM has an excellent ability to capture the spatial patterns of empirical alpha and beta frequency bands, requiring only 5-10 graph eigenmodes to do so. To our knowledge, the presented SGM and M-SGM are the only models currently available that can simultaneously predict both the regional spectra as well as the spatial distribution of the empirical alpha and beta band activity. In the latter task there was no difference in performance between M-SGM and the original SGM. We see that quantitatively, the modified SGM’s modeled spectra captures the MEG spectra better than the original SGM, but its ability to predict the spatial distribution of the empirical alpha and beta frequency bands is no different from that of the original SGM. Please recall, the parameter optimization was done to maximize spectral fits only. A potential future study will involve incorporating both spectral as well as spatial correlations to obtain optimal model parameters. Even though the quantitative performance is only better when comparing spectral correlations, it is to be noted that this modified local mesoscopic model is biophysically similar in formulation to the nonlinear neural mass models.

### 4.1 No natural ordering of eigenmodes

Surprisingly, in our analysis of which graph eigenmodes most contribute to the summation in the M-SGM (Fig. 2C and D and Fig. 3C and D), we found that the eigenmodes that maximally contribute to the spatial patterning of alpha or beta power are not necessarily those that appear as the first few eigenmodes as ordered by their eigenvalue. Although our results clearly indicate that the first few eigenmodes have the highest frequency of participation, almost all eigenmodes exhibit some ability to participate in the model summation. That the “natural ordering” of eigenmodes (i.e. in increasing order of eigenvalue) is not reflected in fitted M-SGM, at least as far as predicting band-limited spatial patterns, is intriguing. Previous eigendecomposition models on graphs have found a strong natural ordering. E.g. Abdelnour et al. report that only the first few Laplacian eigenmodes (with smallest eigenvalues) are required to resemble empirical functional connectivity of resting state BOLD fMRI [5, 41]; similar results were reported by Atasoy and others [42–44]. Other studies involve a series expansion of the graph adjacency or Laplacian matrices [45–49], which also amount to weighting the first natural eigenvectors more heavily in the summation. This discrepancy might reflect the different context here: MEG rather than fMRI, and modeling of spectral power instead of functional connectivity. It is also possible that since our M-SGM model has far more expressability and a richer repertoire of wideband activity spectrum, the model parameters are capable of tuning a wider range of eigenmodes. This aspect was also noted in a recent thorough exposition of complex Laplacian eigenmodes [44]. The ability of the brain’s connectome to engage a wide range of eigenmodes might be an important feature of realistic and rich brain behaviour - an aspect that merits a deeper future exploration.

### 4.2 Limitations, potential applications, and future work

By design, the model’s frequency spectra can exhibit two peaks at most, whereas empirical MEG spectra (see Fig. 1A) have higher frequency peaks than alpha and beta. Although it is possible to generate those higher beta or gamma peak as a primary peak by varying the local time constants, additional peaks cannot be generated using the current model under plausible parameter regimes. In the future, we will extend the model to capture the peaks in the higher beta and gamma regions as well - this will require additional local population coupling and complexity. However, we note that our current model is already fully capable of reproducing the secular frequency fall-off seen in empirical spectra, e.g. the hypothesized 1/*f^α^* behavior.

This model can be extended to investigate various aspects of functional brain activity. Firstly, this work can be extended to investigate temporal state changes in the functional activity. One way would be by introducing a temporal component to the model parameters. Secondly, this work can also be extended to investigate functional activity in different brain states such as sleep, task-focused, and unconscious. As mentioned earlier, we also plan to extend this model to generate peaks in the higher beta and gamma regions of the modeled spectra. Future work will also involve ensuring that the system is stable with the estimated parameters, using adaptive gain control.

This modeling approach can be used to investigate underlying biophysical mechanisms involved in different neurological diseases, as well. For example, we recently estimated regionally varying local model parameters for empirical MEG spectra collected for healthy and Alzheimer’s disease subjects. We then found that the excitatory and inhibitory parameters were differentially distributed for the healthy versus the Alzheimer’s disease subjects, indicating an excitatory/inhibitory imbalance (*submitted*). Such investigations can be extended to various other neurological diseases as well. Lastly, it will be of interest to extend this model to include other modalities such as fMRI and EEG, as well. In these biomedical applications, the parsimony and inferrability of the SGM model will prove to be critical assets, especially in comparison with current high-dimensional and non-linear models.

## Data availability

The code and processed datasets (processed connectivity and distance matrices, and the MEG spectra) used in this work are available at https://github.com/Raj-Lab-UCSF/spectrome-revisited and are based on the original SGM repository [35].

## Acknowledgments

This work was supported by NIH grants R01NS092802/ 183412 and RF1AG062196. The template HCP connectome used in the preparation of this work were obtained from the MGH-USC Human Connectome Project (HCP) database (https://ida.loni.usc.edu/login.jsp). The HCP project is supported by the National Institute of Dental and Craniofacial Research (NIDCR), the National Institute of Mental Health (NIMH) and the National Institute of Neurological Disorders and Stroke (NINDS). Collectively, the HCP is the result of efforts of co-investigators from the University of Southern California, Martinos Center at Massachusetts General Hospital (MGH), Washington University, and the University of Minnesota. Additionally, we would like to acknowledge Kamalini G. Ranasinghe for helping with generating the brain surface renderings.

